# Sodium sulfite as a switching agent for single molecule based super-resolution optical microscopy

**DOI:** 10.1101/2021.10.11.462112

**Authors:** Anders Kokkvoll Engdahl, Oleg Grauberger, Mark Schüttpelz, Thomas Huser

**Affiliations:** Biomolecular Photonics, Bielefeld University, Universitätsstrasse 25, 33615 Bielefeld, Germany

**Keywords:** dSTORM, SOFI, imaging buffers, sulfite, SMLM, localization microscopy

## Abstract

Photoinduced off-switching of organic fluorophores is routinely used in super-resolution microscopy to separate and localize single fluorescent molecules, but the method typically relies on the use of complex imaging buffers. The most common buffers use primary thiols to reversibly reduce excited fluorophores to a non-fluorescent dark state, but these thiols have a limited shelf life and additionally require high illumination intensities in order to efficiently switch the emission of fluorophores. Recently a high-index, thiol-containing imaging buffer emerged which used sodium sulfite as an oxygen scavenger, but the switching properties of sulfite was not reported on. Here, we show that sodium sulfite in common buffer solutions reacts with fluorescent dyes, such as Alexa Fluor 647 and Alexa Fluor 488 under low to medium intensity illumination to form a semi-stable dark state. The duration of this dark state can be tuned by adding glycerol to the buffer. This simplifies the realization of different super-resolution microscopy modalities such as direct Stochastic Reconstruction Microscopy (dSTORM) and Super-resolution Optical Fluctuation Microscopy (SOFI). We characterize sulfite as a switching agent and compare it to the two most common switching agents by imaging cytoskeleton structures such as microtubules and the actin cytoskeleton in human osteosarcoma cells.

## Introduction

Single-molecule localization microscopy (SMLM) utilizes the various photochemical properties of fluorescent dyes to identify different dyes by differences in their intermittent fluorescence emission, thereby allowing for their localization.^1^ In *direct Stochastic Optical Reconstruction Microscopy* (dSTORM), organic fluorophores are subjected to high illumination intensities in an imaging buffer containing primary thiols and enzymatic oxygen scavengers.^2,3^ The thiols react with triplet-state fluorophores to form a semi-stable, non-fluorescent dark state. The dSTORM technique relies upon the use of an enzymatic scavenger system consisting of the enzymes *glucose oxidase* and *catalase*, that when used in combination with a high glucose content in the buffer, will lower the oxygen concentration of the buffer, thereby minimizing the chance of irreversible photodamage to the dye. Either *mercaptoethylamine* (also known as cysteamine and its abbreviation MEA) or *beta-mercaptoethanol* (BME) is typically used to reduce the triplet state of the dye molecule to either the first or second reduced state, so that the dye spends a longer time in the non-fluorescent state. In order to achieve a large fraction of dyes in the off-state, dSTORM relies on illumination intensities on the order of 1 kW cm^-2^ or higher. At these rather high laser intensities, photodamage to the fluorophores is still often the outcome after an extended measurement. In addition, this buffer quickly loses its switching properties (“freshness”), since the enzymatic depletion of oxygen gradually acidifies the buffer. This means that fresh dSTORM buffers typically have a usefulness of a couple of hours, though sealing off the chamber to ambient air can slow down the acidification process. dSTORM buffers are typically mixed directly before use by thawing stock solutions of enzymes, glucose, and thiols, and adding these together. The stock solutions, kept in freezers at -20°C, also have to be remade every few months, as cysteamine is especially prone to decay over time. The other primary thiol used for dSTORM buffers, BME, is a toxic and corrosive chemical with a foul odour that makes working with dSTORM buffers in an unventilated room unpleasant and a health hazard.

Furthermore, a challenge for dSTORM has been to develop high refractive-index imaging buffers. Certain solutions to this problem do exist, such as Vectashield, and the more recent glycerol-glucose buffer.^4,5^ However, in our hands, these buffers need a much higher illumination power in order to properly switch fluorophores to the dark state, than in their aqueous forms. Recently, a high refractive-index buffer containing 90% (v/v) glycerol, sodium sulfite (Na_2_SO_3_) as a chemical oxygen scavenger, and cysteamine as a switching agent, was reported by Hartwich et al^6^. The buffer was superior to ordinary dSTORM buffers in suppressing background signal, a feat the authors explained to be due to an increased efficiency in cysteamine-induced switching in the oxygen-low solution. According to the authors, sulfite was not found to reduce the fluorophore Alexa Fluor 647 (AF647) and switch it to the dark state. Contrary to Hartwich et al., we tested sodium sulfite in aqueous and glycerol-rich buffers and found it to be an even stronger switching agent than cysteamine and beta-mercaptoethanol.

## Results

We performed dSTORM experiments on human osteosarcoma cells (U2OS), where micro-tubulin was labeled with AF647-conjugated secondary antibodies, using a solution of phosphate-buffered saline, 50 mM sodium sulfite, and different concentrations of glycerol (**Figure 1**). At 0% (v/v) glycerol, sodium sulfite caused AF647 to fluctuate its emission brightness, enabling *Super resolution optical fluctuation* (SOFI) microscopy^7,8^, as seen in **Figure 1a**. With the addition of 1% (v/v) glycerol or more, and imaging at illumination intensities of 0.2 kW cm^-2^, AF647 transitioned to the dark state almost instantly, which enabled dSTORM at low laser power (**Figure 1b**, mean frame intensities shown in **FIgure 1d**). **Figure 1e** shows a reconstruction of densely labelled tubulin in U2OS cells imaged for 10,000 frames at 40 ms exposure time using a 647 nm laser in epifluorescence at 0.3 kW cm^-2^. We measured a resolution of approximately 30 nm using Fourier ring correlation.^9^ The experiment was confirmed to work on a setup using a cost-efficient CMOS camera and even lower intensities of ∼0.2 kW cm^-2^ (**Figure S1**).

**Figure 1:**
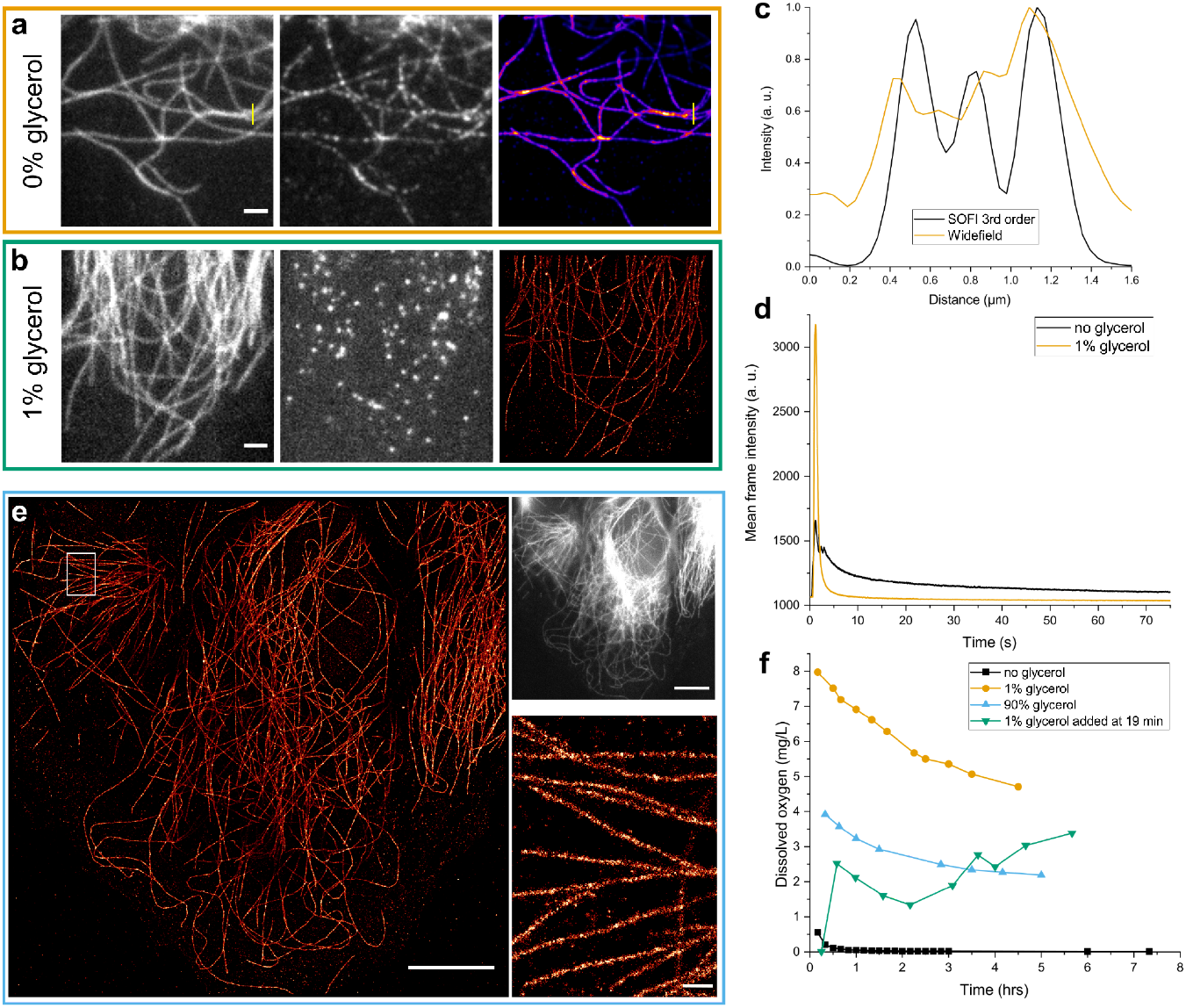
A simple solution of PBS containing 50 mM sodium sulfite can cause AF647 to switch from stable fluorescence to intermittent fluorescence emission, enabling 3^rd^ order SOFI reconstruction, as shown in **a**, where 4000 frames were used for the SOFI calculation. **(b)** The addition of 1% (v/v) glycerol to the buffer makes sulfite an efficient switching agent for AF647, enabling single molecule localization. **(c)** A line profile through AF647-labeled tubulin in panel **a** shows the resolution improvement of 3rd order SOFI. **(d)** Mean frame intensities for measurements **a** and **b** show how quickly off-switching occurs at 1% (v/v) glycerol as compared to when no glycerol is present. **(e)** dSTORM reconstruction of 10,000 frames, recorded at 0.3 kW cm^-2^ of 647 nm laser power, using 40 ms exposure per frame. Top right inset shows the widefield diffraction-limited image. The inset in the reconstruction is shown in the bottom right corner. **(f)** Measurement of dissolved oxygen in various buffers shows how the addition of glycerol quenches the oxygen removal effect of sodium sulfite. Scale bars: **a**,**b**: 2 µm. **e**: 10 µm (left image and top right image), 0.5 µm (bottom right image).

In order to better understand how the addition of glycerol affects the buffers, we measured the concentration of dissolved oxygen using an oxygen meter (O2 Transmitter 170 air, Mettler Toledo, Switzerland). The results are shown in **Figure 1f**. As expected, sodium sulfite breaks down dissolved oxygen in an aqueous solution. However, the addition of 1% glycerol disrupted the process, so that instead of an oxygen concentration nearing the detection limit, over the course of multiple hours it approached a 40-50% saturated concentration. This raises the question of whether or not the “switching” seen in the glycerol-containing sulfite buffers is due to fluorophores being reduced to the off-state, or simply photobleaching by oxygen. However, we conclude that the switching reaction is reversible, as the observed “blinking” is very different from the permanent fade-away of fluorophores that occurs in a solution containing merely PBS and glycerol. Another possibility is triplet-state-induced blinking by molecular oxygen, but the clear facilitation of this process in the sulfite buffer, as compared to in normal PBS, makes it clear that sulfite should be the main source of blinking.

In order to test the performance of sulfite as a switching agent, we performed the initial switch-off phase for AF647-stained tubulin in U2OS cells at 0.3 kW cm^-2^ for oxygen scavenging buffers containing either 50 mM of MEA, 100 mM of BME, or 50 mM of SO3, the two first being common concentrations used in dSTORM microscopy. As our oxygen scavenging buffer also contained approx. 7% (v/v) glycerol from the prepared stock solutions, no additional glycerol was added. **Figure 2a** shows time lapse images taken in experiments with the three different switching agents, where the sulfite-containing buffer is clearly the first to enter the single-molecule regime, although slower than in the “simple sulfite” buffer shown in **Figure 1d**, and with a much lower overall intensity regime, as seen in **Figure 2b**. The initial frames in a dSTORM experiment, that are usually filtered out in the reconstruction phase due to the presence of too many emitting fluorophores in a single frame, were therefore fewer, and the analysis could be started sooner. Fits of the mean intensity of frames over time showed that off-switching of emitters occurred faster, as seen in **Figure 2c**. We found that the sulfite-containing buffer on average resulted in a decay time constant for the switch-off phase of AF647 of 2.6 ± 1.3 s, compared to 3.8 ± 0.4 s, and 4.9 ± 1.0 s for MEA- and BME-containing dSTORM buffers. Using SOFIevaluator on the data^10^, showed that on-times are lower for emitters when using sulfite, than at corresponding concentrations of thiol-based agents (**Figure 2d,e**). When comparing the switching agents at a laser power of 0.7 kW cm^-2^, we found that sulfite on average resulted in an emitter intensity of 70% of that obtained when using cysteamine, and only 50% of the intensity when using beta-mercaptoethanol (**Figure 2f**). However, since the offset is also much lower in the sulfite switching buffer than that for BME, the localization precision is similar for the two cases (14.5 nm for sulfite, 15.5 for BME, **Figure 2g**). The number of unique localizations (i.e. corrected for emitters that are on for more than one frame) per frame and area is similar for all three switching agents (**Figure 2i**). In general, cysteamine performed best for dSTORM at 0.7 kW cm^-2^, with a localization precision of 12.8 nm.

**Figure 2.**
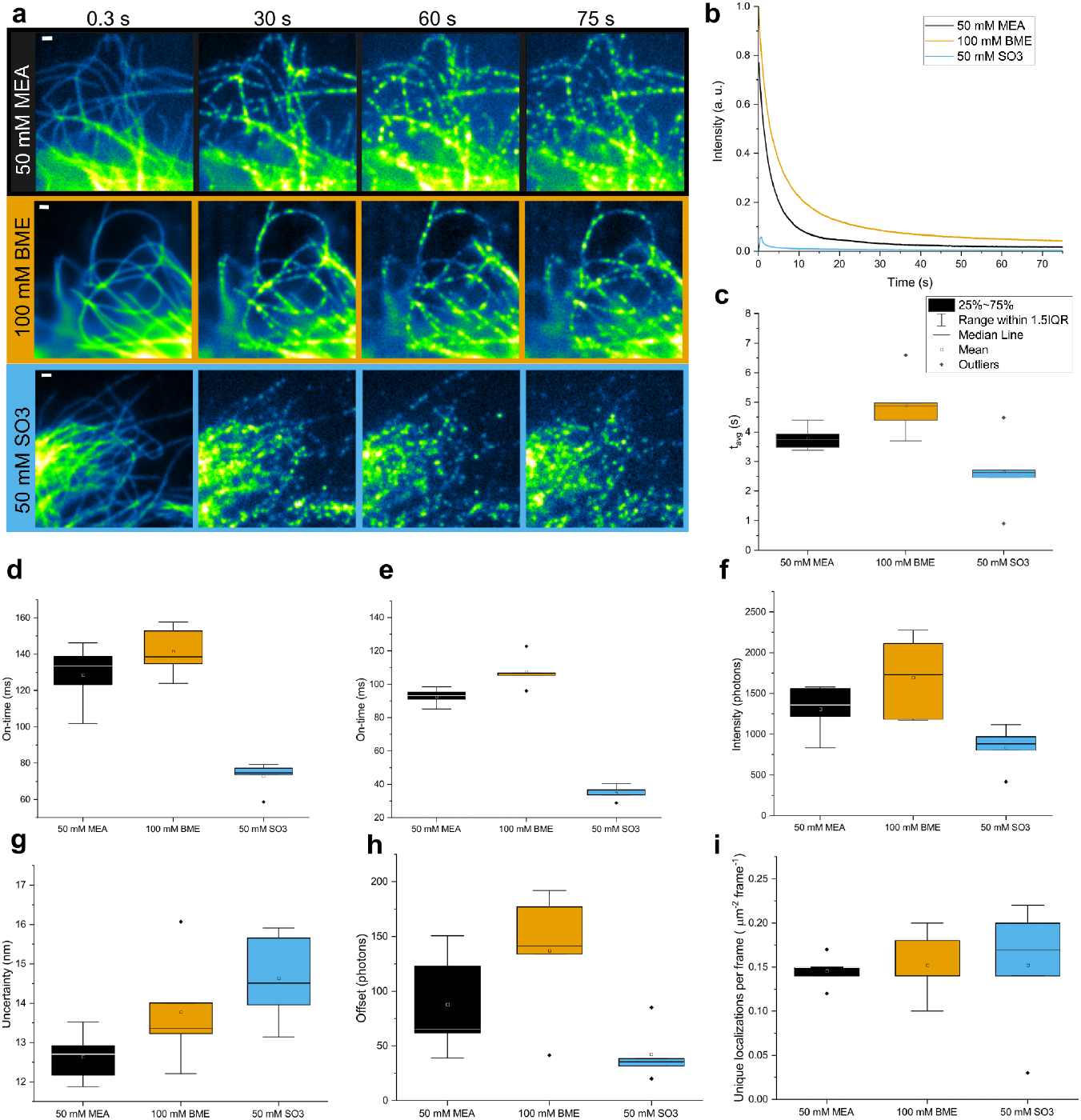
Performance of sodium sulfite in an enzymatic oxygen scavenging buffer, as compared to cysteamine and beta-mercaptoethanol. **(a)** Time-lapse of AF647-stained microtubules in U2OS cells, when subjected to 0.3 kW cm^-2^ of 647 nm laser light, using the three switching agents. **(b)** Mean frame intensities during the experiments, relative to the highest value. **(c)** Intensity decay constants for MEA, BME and SO3. **(d**,**e)** On-time of AF647 fluorophores when subjected to 0.3 kW cm^-2^ or 0.7 kW cm^-2^ of laser power, as calculated by SOFIevaluator. **(f)** Intensity of localizations at 0.7 kW cm^-2^ calculated by ThunderSTORM.^16^ **(g)** Localization uncertainty of **f. (i)** Unique localizations per frame and µm^-2^.

Higher index STORM buffers are those where sulfite outperforms thiols as a switching agent. We devised an enzymatic oxygen scavenging buffer containing roughly 70% (v/v) glycerol and 50 mM sodium sulfite. Off-switching of AF647-conjugated antibodies occurred much faster in 70% glycerol, as opposed to the standard sulfite dSTORM buffer containing 7% glycerol, as seen in **Figure 3b**. In 70% glycerol, the initial off-switching phase was finished in a matter of seconds at 0.3 kW cm^-2^ of 647 nm wavelength laser exposure, whereas 90 seconds of laser exposure was insufficient to turn a significant number of emitters to the dark state in the low-glycerol buffer. **Figure 3a** shows a dSTORM reconstruction of microtubules in U2OS cells stained with AF647-conjugated antibodies in the 70% glycerol-containing sulfite buffer using 5000 frames at this low laser power density using 15 ms exposure time per frame. A drawback of the high glycerol concentration is the loss of brightness of localized emitters, where only approx. 400 photons per localization per frame are detected, vs. approx 900 photons for the 7% glycerol buffer (**Figure 3c**). This drawback is, however, somewhat reduced by the much lower background signal for localizations in the high glycerol buffer (**Figure 3d**). The localization uncertainty is higher in 70%, than in 7% glycerol, with approx. 16.2 nm, vs. 14.5 nm, respectively. Interestingly, the localization uncertainties in 70% glycerol buffers are similar for laser illuminations of 0.3 kW cm^-2^ and 0.7 kW cm^-2^, although with a higher spread across measurements. This indicates that the “on-time” of AF647 on average occurs on a shorter time frame than the 15 ms camera exposure time. The drawback of imaging at 0.3 kW cm^-2^ illumination intensity is that a greater amount of emitters are in the on-state at any given time, and therefore more emitters had to be filtered out in our analysis (**Figure 3f**). The 0.3 kW cm^-2^ laser power in a high glycerol concentration sulfite buffer can be a viable alternative when higher laser powers or imaging time are limited, as the localization uncertainty is not much higher than that used for 0.7 kW cm^-2^ power density in the standard dSTORM buffer. On average, more multi-emitter events were recorded in the lower glycerol concentration buffers than in the higher, as seen in **Figure 3f**. In our experience, a dSTORM buffer containing 70% glycerol reduces the switching efficiency of cysteamine and beta-mercaptoethanol as compared to aqueous solutions.

**Figure 3:**
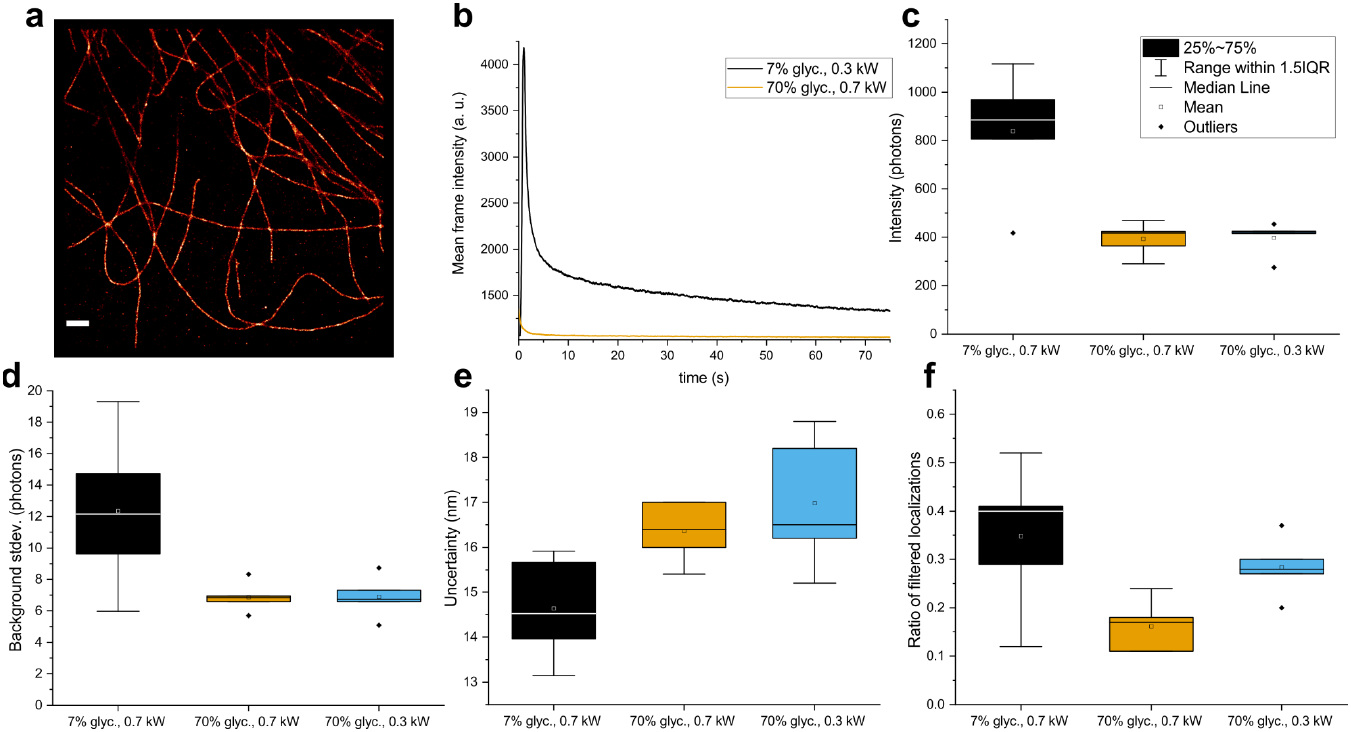
Effect of a high glycerol concentration in the oxygen scavenging dSTORM buffer with sodium sulfite as a switching agent. **(a)** Image reconstruction of 5000 frames with 15 ms exposure at 0.3 kW cm^-2^ laser power using a buffer containing 70% glycerol and 50 mM sodium sulfite. **(b)** Off-switching duration of AF647 in 50 mM sulfite in 70% glycerol is nearly instant upon laser radiation, as compared to 50 mM sulfite in 7% glycerol. **(c)** Box plot of intensities of localized emitters for AF647 using 50 mM sulfite in 7% glycerol at 0.7 kW cm^-2^, 70% glycerol at 0.3 kW cm^-2^, and 70% glycerol at 0.7 kW cm^-2^, respectively. **(d)-(f)** Background standard deviation, localization uncertainty and the ratio of localizations that were filtered out, respectively. Data sets are the same as for **c**. Scale bar for panel **a**: 1 µm.

### Switching of other fluorophores

The fluorophore Alexa Fluor 488 (AF488) behaves similarly to AF647 when subjected to a sulfite-containing buffer, where emission fluctuations that enable SOFI reconstructions are observed when AF488 is exposed to a glycerol-free solution. When 10% (v/v) glycerol is added to the solution, we observe off-switching of dyes that enables dSTORM at relatively low illumination intensities. **Figure 4** displays SOFI and dSTORM experiments of actin in U2OS cells stained with Phalloidin-AF488 using a sulfite buffer without any enzymatic oxygen scavenging system. We performed SOFI of 4000 frames (**Figure 4b**), imaged at 15 ms frame exposure using 0.6 kW cm^-2^ of 488 nm laser power in a buffer containing 50 mM sulfite in PBS. Using SOFIevaluator to analyse the data,^10^ we obtained approx. 25% useful SOFI signal (**Figure 4f**) and a clear separation of SOFI signal from noise (**Figure 4g**). Switching on-time (tau) was measured to approx. 5 ms for 0.6 kW cm^-2^, indicating that an improvement of these data is still possible. **Figure 4h** and its inset **Figure 4i** show the widefield image (on the left) and the dSTORM reconstruction of 30,000 frames of F-actin labelled with AF488 when exposed to 0.5 kW cm^-2^ of 488 nm radiation. Here, we used a UV laser (378 nm) to reactivate fluorophores in the dark state, and obtained a 29 nm resolution, indicating that fluorophores in this simple imaging buffer are switched off reversibly, and do not easily photobleach.

**Figure 4:**
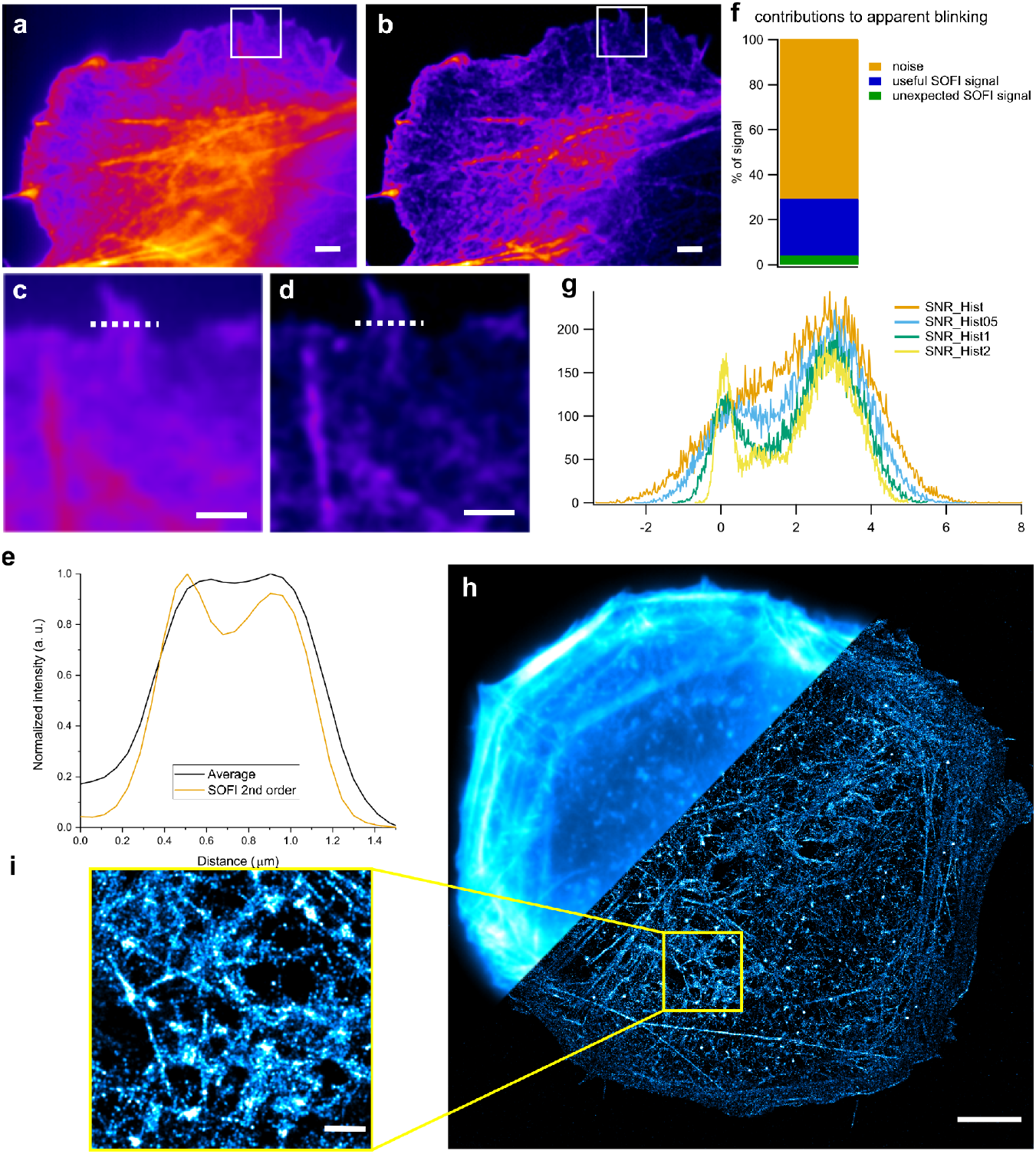
Switching behaviour of AF488 demonstrated by super-resolution microscopy of U2OS cells. Here, the actin cytoskeleton was stained with Phalloidin-AF488 and imaged using a sulfite buffer without an enzymatic oxygen scavenging system. **(a), (b)**, Average intensity and 2^nd^ order SOFI of 4000 frames, respectively. **(c), (d)** Insets from **a** and **b**, with line plots of the dashed line shown in **e. (f), (g)** Ratio of useful SOFI signal to noise and unaccounted SOFI signal, and signal-to-noise histograms of the image data, respectively. The data were analysed using SOFIevaluator. **(h)** Left: widefield image and right: dSTORM reconstruction of F-actin in U2OS cells stained with phalloidin-AF488. **(i)** Inset of reconstruction **h**, with an FRC resolution of approx. 30 nm. Scale bars: **a**,**b**: 2 µm. **c**,**d**: 1 µm. **h**: 5 µm. **i**: 1 µm.

We also tested the dyes AF 568 and CF 568 with the sulfite buffer, but did not observe any notable dye blinking.

## Conclusion

We conclude that sodium sulfite, together with glycerol, leads to intermittent fluorescence emission of select Alexa Fluor dyes, AF488 and AF647. The fact that sulfite also switches AF488 distinguishes it from the phosphine *tris(2-carboxyethyl)phosphine* (TCEP), which only acts as a switching agent for AF647.^11^ Since sodium sulfite is also an oxygen scavenger, it can be used in a simple dSTORM buffer containing only PBS and low concentrations of glycerol, although adding glycerol significantly lowers its oxygen scavenging potential. When used without glycerol, it acts as an agent that causes fluctuations in dye brightness well suited for SOFI image reconstructions. It is possible that the main role of glycerol is to prevent oxygen from being removed by sulfite, and that oxygen in some concentration is needed for the most efficient switching by sulfite. The efficiency of sodium sulfite as a switching agent is highest without the enzymatic oxygen scavenging buffer, or in a buffer with a high glycerol concentration. In an oxygen scavenging buffer, sulfite is similar to MEA and BME when imaging at lower illumination intensities (P < 0.5 kW cm^-2^), with especially its efficacy for off-switching and low on-times of AF647 marking it exceptional. In our experience, sodium sulfite works best in a high glycerol-concentration dSTORM buffer, outshining cysteamine as the switching agent. The mechanism of switching, and glycerol’s role thereof, is not known. Another unknown is the cause of the stark reduction in brightness of the fluorophore AF647 in the sulfite switching buffer, similar to what has recently been reported of the Vectashield dSTORM buffer^12^. Once the mechanism behind this behavior is established, it might be possible to find a switching buffer that switches the fluorophore emission as well as sulfite, while maintaining the brightness observed with the primary thiols.

The potential applications of sulfite as a switching agent could be many. In the recent years we have seen a surge in emerging low-cost equipment for (super-resolution) microscopy,^13,14,15^ but dSTORM still relies on rather expensive chemical reagents that also require access to a -20°C freezer, such as primary thiols and the enzymes for the oxygen scavenging system. Recording good SOFI experiments can also be difficult. Our simplest sulfite buffers, i.e. 50 mM sulfite in aqueous phosphate-buffered saline with the optional addition of glycerol for dSTORM, simplifies experiments significantly. More importantly, it lowers the traditional power threshold for dSTORM, which can be useful on many setups that aren’t equipped with powerful lasers. We hope that these buffers can be useful to the microscopy community and further spread the applications of localization and fluctuation microscopy.

**Table 1:** Summary of the results for different buffers containing an enzymatic oxygen scavenger system used for switching of AF647. Mean values of five different experiments per buffer are summed up here, with standard deviation in parentheses.

| Buffer composition | Laser power (kW cm<sup>-2</sup>) | Switch-off-decay constant (at 0.3 kW cm<sup>-2</sup>) (s) | On-time (ms) | Intensity (photons) | Loc. uncertainty (nm) | Offset (photons) | Ratio of filtered localizations | Unique localizations (locs. frame<sup>-1</sup> μm<sup>-2</sup>) |
| --- | --- | --- | --- | --- | --- | --- | --- | --- |
| 50 mM MEA | 0.7 | 3.8(0.4) | 92.6(5.0) | 1310(305) | 12.6(0.6) | 87.9(46.8) | 0.28(0.1) | 0.15(0.02) |
| 100 mM BME | 0.7 | 4.9(1.1) | 107.4(9.6) | 1697(511) | 13.8(1.4) | 137.2(58.7) | 0.41(0.13) | 0.15(0.04) |
| 50 mM SO<sub>3</sub>, 7% glycerol | 0.7 | 2.6(1.3) | 34.8(4.3) | 839(262) | 14.6(1.2) | 42.3(25) | 0.35(0.15) | 0.15(0.07) |
| 50mM SO<sub>3</sub>, 70% glycerol | 0.7 |  | 38.5(6.7) | 393(69) | 16.4(0.7) | 23.9(3.8) | 0.16(0.05) | 0.16(0.05) |
| 50 mM SO<sub>3</sub>, 70% glycerol | 0.3 | 4.3(0.8) | 60.0(4.7) | 398(70) | 17.0(1.5) | 20.9(5.8) | 0.28(0.06) | 0.17(0.04) |

