## Supporting information for "Sodium sulfite as a switching agent for single molecule based super-resolution optical microscopy"

#### Mixing sodium sulfite buffers

Sodium sulfite (CAS nr: 7757-83-7) can be stored in Eppendorf tubes at room temperature in concentrations of 1-2 M in bidest water for several weeks, but should for the best results be mixed fresh from powder before experiments.

Phosphate-buffered saline (PBS) was mixed in 50 mL Falcon tubes by adding 2 tablets to 40 mL of bidest water. This gives a concentration of 10x PBS, which could then be diluted to working concentrations.

Glycerol (ROTIPURAN, 99.5%, ROTH) was mixed to stock solutions of 10% v/v in bidest water, or added as pure reagent to the high glycerol-concentration buffers.

##### Enzymatic oxygen scavenger dSTORM buffers

Stock solutions of enzymes, glucose and MEA were prepared according to van de Linde (2011)[1] and stored at -20C until used. The finished dSTORM buffer consisted of 0.001 mg/mL catalase (C30, Sigma Aldrich), 0.05 mg/mL glucose oxidase (G2133, Sigma Aldrich), 0.04 g/mL glucose, and either 50 mM MEA, 100 mM BME or 50 mM sulfite. Additionally, the buffers also contained 0.2 µM TCEP (Sigma Aldrich), 1.25 µM KCl, 1 µM Tris-HCl (pH=7.5) and approx. 7% (v/v) glycerol. The buffers were mixed with 10x PBS so as to contain approx. 1x PBS at the end concentration.

High glucose oxygen scavenging buffers contained approx. 70% (v/v) glycerol, and 0.02 g/mL glucose.

##### Measurements of dissolved oxygen in imaging buffers

The buffers shown in **FIgure 1** were mixed in 50 mL Falcon tubes to end volumes of 10 mL each. The pO2 sensor was calibrated in bidest water with constant air being bubbled through, until it settled. Then it was placed in the desired buffer solution, and the measurement began. Each time series was measured separately.

#### Preparation of cell samples

Human osteosarcoma cell culture was seeded in Ibidi 8-well plates and grown in DMEM for at least 24 hours. Fixation was done according to Jimenez et al.,[2] using 0.5% glutaraldehyde in a cytoskeleton-preserving buffer containing 0.25% Triton X-100, following extraction with 0.1% glutaraldehyde in cytoskeleton-preserving buffer. Cell samples were washed three times with PBS, then glutaraldehyde-induced autofluorescence was bleached using 0.1% sodium borohydride (NaBH4) in PBS. Blocking was performed either with 5% Bovine serum albumin (BSA) and 0.1% Triton X-100 in PBS or using 0.22% of gelatine in PBS, and left on the sample for at least one hour. Tubulin staining was performed using mouse anti-alpha-tubulin IgG (T5168 or T6199, Sigma Aldrich) and mouse anti-beta-tubulin IgG (T8328, Sigma Aldrich) antibodies, each at 1:300 dilution in 1% BSA, 0.02% Triton X-100 in PBS overnight at 4C, or the same antibodies in 0.22% gelatine with 0.02% Triton X-100. As a secondary antibody, goat anti-mouse conjugated with Alexa Fluor 647 (A-21237, Invitrogen) at 1:200 dilution in 0.1% BSA with 0.02% Triton X-100 in PBS, or alternatively, 0.22% gelatine with 0.02% Triton X-100 in PBS, was used. Staining was performed at room temperature for 90 minutes. Following washing in PBS, samples were postfixated using 0.1% paraformaldehyde in PBS for 3 minutes. Staining of actin filaments was done using Phalloidin-AF488 (A12379, Invitrogen) in a 1:20 dilution in 1% BSA in PBS for 60 minutes at room temperature.

### Microscope setups

#### Acquisitions with an EMCCD camera-setup

We used a custom-modified Olympus IX71 microscope body with optionally a IX2-NPS nosepiece. We used an Andor iXon DV887DCS-BV 512 x 512 16 um pixel EMCCD camera, with, after magnification, a pixel size that measured 0.113 µm. The laser source was an Argon-Krypton-Ion laser (Innova 70C, Coherent) with laser lines at 488 nm and 647 nm, and for photoactivation of fluorophores, a UV laser with a wavelength of 378 nm, and 16 mW (Coherent Cube). An acousto-optic tunable filter (AOTF) (AOTRFnC-VIS-TN 1001, AA Opto Electronic) was used to select wavelengths and control laser output. The objective lens was an Olympus Apo N 1.49 TIRF. The excitation filter was a multi-line bandpass filter (497/655 H, F58-200, AHF Analysentechnik AG). Emission filters were a long pass (Razor Edge 647, Semrock) and a bandpass (ET700/75 M, Chroma), for the 647 nm line, and a long pass (Edge Basic 488, Semrock) and band pass (520/40ET, Chroma) for the 488 nm laser line.

Acquisitions are listed in the table below:

**Table S1:** Image acquisition details. All acquisitions were recorded with the laser in epi-fluorescence mode.

| **Figure** | **Number of frames used for reconstruction** | **Exposure time (ms)** | **Laser line (nm)** | **Laser power (kW $cm^{-2}$, approximate)** | **Dye** |
|---|---|---|---|---|---|
| 1a | 4,000 | 15 | 647 | 0.3 | AF647 |
| 1b | 5,000 | 15 | 647 | 0.3 | AF647 |
| 1e | 10,000 | 40 | 647 | 0.3 | AF647 |
| 2 | 5,000 | 15 | 647 | 0.3, subsequent 0.7 | AF647 |
| 3a | 5,000 | 15 | 647 | 0.3, subsequent 0.7 | AF647 |
| 4a,b | 4,000 | 15 | 488 | 0.6 | AF488 |

| | | | | | |
|---|---|---|---|---|---|
| 4h,i | 30,000 | 50 | 488, 378 | 0.5, <0.01 | AF488 |
| S1 | 10,000 | 50 | 640 | 0.2 | AF647 |

Single-molecule localization data were fitted using the ThunderSTORM plugin,[3] using the following camera settings: Pixel size: 113 nm. Photoelectrons per ADU count: 20.94 Base level: 1014. Gain: 54.61. For the rest of the analyses, the localizations were filtered with these values: uncertainty < 100 & intensity < 8000 & sigma > 67.56 & sigma < 184.5. SOFI data were analysed using Localizer or analysed using SOFIevaluator.[4,5]

#### Buffer comparison experiments

For each buffer, five measurements at separate ROIs were made at two illumination intensities, first at 0.3 kW $cm^{-2}$, then at 0.7 kW $cm^{-2}$ for the same ROI. After localization analysis, the median values of these measurements were then used to construct the box plots seen in **Figure 2** and **Figure 3**.

#### Acquisitions on a low-cost setup

A previously described low-cost modular dSTORM microscope was used to display that the buffers can work for CMOS cameras. The laser source was a Toptica iBeam Smart 640 with 150 mW max output. The filters that were used were a ZET 640/10 clean-up, Chroma, and a RazorEdge 647, Semrock BLP01-647R-25 for the detection path. The objective lens was an Olympus Apo N 1.49 TIRF objective, used together with a tube lens with 100 mm focal length. The camera was a IDS uEye UI-3270CP-M-GL Rev. 2 with a pixel size of 3.45 um before magnification. Camera offset was 134.6 in gray values, gain 0.162 ADU/$e^-$, Quantum efficiency: 0.53. Pixel size after magnification was measured to 0.105 µm. We used Micromanager 2.0 for the data acquisition.[6] An example dSTORM experiment on this setup is shown in **Figure S1**, where we used 50 mM sulfite in 1% glycerol-containing PBS and approx. 0.2 kW $cm^{-2}$ of 640 nm sample illumination.

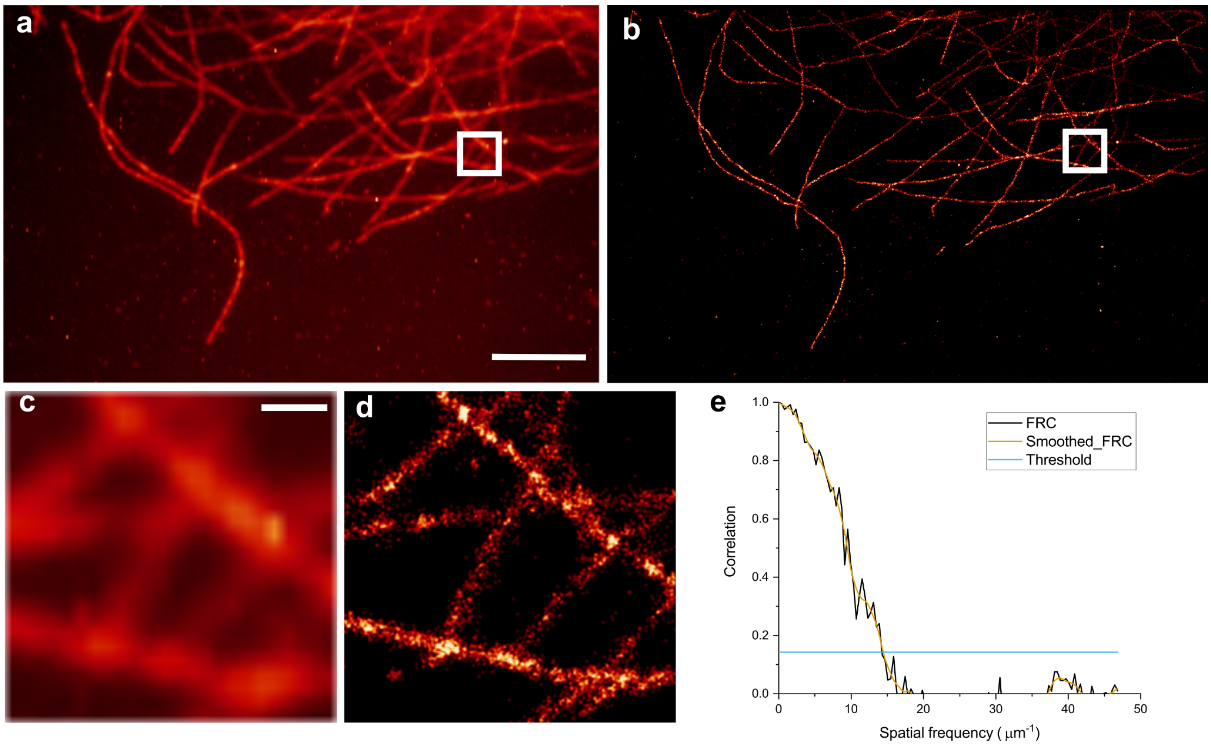

**Figure S1:** dSTORM imaging of AF647-stained tubulin on a low-cost setup using a CMOS camera and a 50 mM sulfite buffer with 1% glycerol.. **a**: Standard deviation of 10,000 frames. **b**: Reconstruction of 10,000 frames. **c**: Inset from **a**. **d**: Inset from **b**. **e**: Fourier-ring correlation of **d**, revealing a 69 nm resolution. Scale bars: **a**: 5 µm. **c**: 0.5 µm.


1. van de Linde, S. *Photoswitching of Organic Dyes and Single Molecule Based Super Resolution Imaging*. (2011).
2. About samples, giving examples: Optimized Single Molecule Localization Microscopy. *Methods* **174**, 100–114 (2020).
3. Ovesný, M., Křížek, P., Borkovec, J., Svindrych, Z. & Hagen, G. M. ThunderSTORM: a comprehensive ImageJ plug-in for PALM and STORM data analysis and super-resolution imaging. *Bioinformatics* **30**, 2389–2390 (2014).
4. Dedecker, P., Duwé, S., Neely, R. K. & Zhang, J. Localizer: fast, accurate, open-source, and modular software package for superresolution microscopy. *J. Biomed. Opt.* **17**, 126008 (2012).
5. Moeyaert, B., Vandenberg, W. & Dedecker, P. SOFIevaluator: a strategy for the quantitative quality assessment of SOFI data. *Biomed. Opt. Express* **11**, 636–648 (2020).
6. Edelstein, A. D. *et al.* Advanced methods of microscope control using µManager software. *J Biol Methods* **1**, e10–e10 (2014).
